# Machine learning for randomised controlled trials: identifying treatment effect heterogeneity with strict control of type I error

**DOI:** 10.1101/330795

**Authors:** Chris C. Holmes, James A. Watson

**Affiliations:** Department of Statistics, University of Oxford, Oxford, UK; Nuffield Department of Medicine, University of Oxford, Oxford, UK; Mahidol Oxford Tropical Medicine Research Unit, Mahidol University, Bangkok, Thailand

## Abstract

**Background:** It is widely acknowledged that retrospective exploratory analyses of randomised controlled trials (RCTs) seeking to identify treatment effect heterogeneity (TEH) are prone to bias and false positives. Yet the increasing availability of multiple data modalities on subjects and the desire to learn all we can from trial participants motivates the inclusion of such analyses within RCTs. Coupled to this, widespread advances in AI and machine learning (ML) methods hold great potential to utilise such data to characterise subjects exhibiting heterogeneous treatment response.

**Methods:** We present new learning strategies for RCT ML discovery methods that ensure strict control of the false positive reporting rate at a pre-specified level. Our approach uses randomised data partitioning and statistical or ML based prediction on held-out data. This can test for both crossover and non-crossover TEH. The former is done via a two-sample hypothesis test measuring overall predictive performance of the ML method. The latter is done via ‘stacking’ the ML predictors alongside a classical statistical model to formally test the added benefit of the ML algorithm. An adaptation of recent statistical theory allows for the construction of a valid aggregate p-value. This learning strategy is agnostic to the choice of ML method.

**Results:** We demonstrate our approach with a re-analysis of the SEAQUAMAT trial. We find no evidence for any crossover subgroup who would benefit from a change in treatment from the current standard-of-care, artesunate, but strong evidence for significant noncrossover TEH within the artesunate treatment group. We find that artesunate provides a differential benefit to patients with high numbers of circulating ring stage parasites.

**Conclusions:** Our ML approach combined with the use of computational notebooks and version control can improve the robustness and transparency of RCT exploratory analyses. The methods allow researchers to apply the latest ML techniques safe in the knowledge that any declared associations are statistically significant at a user defined level.

## Introduction

In the medical sciences randomised controlled trials (RCTs) provide the gold standard for evidence evaluation of novel treatments and health interventions. The growing accessibility and recording of data modalities, arising from genetics, medical imaging, genomics, and electronic health records alongside breakthroughs in machine learning (ML) and AI provide opportunities for scientific discovery by characterising patient strata with respect to treatment effect ^1;2^. This can improve patient outcomes and optimise treatment recommendations. However, exploratory analyses of RCTs and correct interpretations of these analyses are difficult ^3;4^ and controversial ^5^. Data analytic tools such as AI algorithms ^6^ are particularly attractive for identifying treatment effect modifiers in RCTs due to their hypothesis free nature and ability to learn by example. While there have been numerous recent papers on technical developments and novel methods for subgroup analysis and treatment effect heterogeneity (TEH) ^7-15^, we know of none to date who have considered ML paradigms which provide strict control of the false positive rate (type I error). Medical statisticians know how to assess the evidence when the subgroups or interactions are predefined and the models are explicit, by counting the “degrees of freedom”, or number of free parameters, in the model and using formal tests of hypotheses ^16-18^. But for ML algorithms the models are designed to adapt their complexity and dependency structures to the underlying problem during the training phase and hence notions of counting parameters become meaningless. The question then remains of how to assess the true evidence of benefit following ML discovery?

We show that it is possible to train ML methods, alongside conventional statistical models, to analyse RCT data and provide a global hypothesis test for the presence of TEH. Our method can formally test the presence of patient subgroups (crossover TEH) and also formally test the added predictive benefit of the ML algorithm by ‘stacking’ the ML predictions alongside predictions from a baseline ‘vanilla’ statistical model. ML algorithms are only justified if their predictive benefit can be proven superior to simpler and more interpretable methods. This framework has important implications for how existing data can be used in a principled manner for trusted hypothesis generation. We hope that our approach will motivate careful *a priori* construction and monitoring of statistical analysis plans utilising the latest ML techniques in RCTs. Such plans are necessary to ensure optimal evidence evaluation and learning through retrospective discovery of TEH.

Our formal approach is illustrated step-by-step via an open source RMarkdown computational notebook ^19^ which uses random forests (RFs) ^20^ to retrospectively analyse a randomised trial in severe malaria ^21^. Throughout this paper we refer to subgroup analysis and TEH interchangeably. Clinically relevant subgroups are a consequence of TEH, in that a subgroup is said to occur when the optimal treatment allocation changes (crosses over), whereas heterogeneity is broader in suggesting any systematic differential in the effectiveness of any one treatment (see panel 1). It is important to distinguish between such crossover and non-crossover events, the former resulting in an optimal treatment allocation that is dependent on patient characteristics ^22^.

## Methods

We support the principle that subgroups of clinical importance identified through a retrospective analysis of trials data, from trials not explicitly designed to identify these subgroups, ultimately need to be validated in a focused, independent, follow-up RCT. Subgroup analysis typically exploits data from trials that were designed to answer a different primary question not involving subgroups and hence the analysis cannot by itself provide a complete picture of the evidence. In this respect, the ML analysis should seek to establish the strength of evidence that any heterogeneous treatment effects are real (true-positives). Establishing and controlling the false-positive rate of the discovery procedure mitigates the risk of following false leads in subsequent confirmatory trials targeting the putative subgroup, and aids in the communication of the evidence from the analysis. The following sections outline a formal methodology for exploratory analysis with strict control of the type I error.

### Pre-defining the ML subgroup statistical analysis plan (ML-SSAP)

Modern statistical and ML methods are able to automate the discovery of subgroups in highdimensional data, and statistical scripting and programming packages such as R, Python, or Stata, allow the analyst to construct routines that take trial data as input and apply statistical or ML models to the data to identify potential TEH. Here we consider both crossover TEH whereby the subgroup is characterised by the set of patients predicted to benefit from a change in treatment compared to the current standard-of-care, and non-crossover TEH whereby the standard-of-care is everywhere optimal but its benefits vary systematically across patient strata. The standard-of-care should be defined prospectively (before looking at the data), even if the analysis is retrospective.

In order to maintain the transparency of the evidence, the ML-SSAP should be specified before any exploration of the primary RCT data has taken place. Failure to do so runs the risk of biasing the results ^23^. When formulating the analysis plan, covering both the ML or statistical method (model) used for discovery, and the set of potential stratifying measurements used by the method, researchers should be cautioned against throwing in every possible variable and every flexible ML method. There is a principle here of ‘no free lunch’, or rather ‘no free power’. The choice of discovery method and the potential variables to include is an important step. Methods that trawl through measurements looking for interactions are not panaceas, nor substitutes for careful thought, and the more judicious the *a priori* data selection and choice of discovery model the higher the expected power and ability of the analysis to detect true effects ^24^.

The analysis plan should also include the specification of a test statistic that can compare overall patient benefit between any two groups and which can be used to quantify the type I error when declaring beneficial subgroups. The form of this test statistic is study specific and should relate to the clinical outcome of interest such as survival time, cure rate, or a quantitative measurement of treatment benefit. This will typically match that used in the original study protocol of the primary trial.

### False-positive control of crossover interactions: subgroup detection

The subgroup analysis refers to the discovery of crossover TEH whereby the optimal treatment allocation changes.

We use a held-out data approach to construct a test for a global null hypothesis of “no true crossover TEH (subgroups)”. Fig 1 illustrates this procedure using the example of a primary two-arm RCT where the original trial failed to detect an overall benefit of the experimental treatment. The approach is as follows. The trial data are repeatedly randomly divided into two subsets, with the ML method fitted independently and separately to each subset. The ML algorithm (or statistical model) is trained on each half of the data separately and is then used to predict the individual treatment effects, and thus the optimal treatments for subjects, in the corresponding other half of ‘held-out’ data. Combining the resulting subjects whose heldout predicted optimal treatment assignment differs from the standard-of-care forms a held-out subgroup of size *n*_*s*_ from the original trial of sample size *n*. The actual treatment administered to these subjects in the primary RCT is random, such that in a balanced two-arm trial we would expect half of the subjects,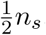, to have received the standard treatment and the other half the experimental treatment. This then facilitates a two-sample hypothesis test, using the test statistic defined in the analysis plan, with a null hypothesis of “no improved subject benefit identified through the ML subgroup analysis plan”. The hypothesis test compares the outcomes of the patients whose optimal treatment was predicted by the ML algorithm to be the experimental treatment and who received the experimental treatment, to those predicted to benefit from the experimental treatment but who received the standard-of-care. A one-sided test would be appropriate if the test statistic measures patient benefit. If there is no true benefit then the resulting p-value is uniformly distributed on [0,1]. If *K* iterations of this procedure are run, randomising the data-split at each iteration, then we obtain corresponding p-values *{p*_1_, *…, p*_*K*_ *}*. We note that these are conservative in that the discovery model on each subset has half the sample size to identify the subgroups. Finally the p-values *{p*_1_, *…, p*_*K*_ *}* can be aggregated to compute a global significance test for the presence of a benefiting subgroup. This aggregation can be done by adapting a method for p-value aggregation in high dimensional regression ^25^. In brief, if *α* is the level of control of the type I error (this is usually set to 0.05), then the set of p-values can be merged into one test using the following formula:

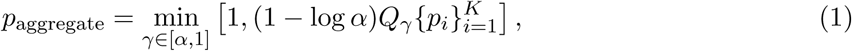

where 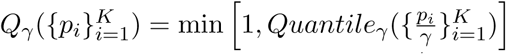. *Quantile*_*γ*_ (*·*) computes the *γ* quantile of the set of p-values which have be scaled by 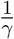. This procedure sweeps over *γ ∈* [*α,* 1] to find the minimum value in *Q*_*γ*_. Alternately the analyst could fix *γ* in the analysis plan, such as *γ* = 0.5 to select the median p-value,

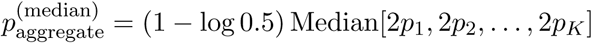

Note, if a true subgroup exists in the population from which the RCT trial participants are drawn, then 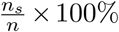 estimates the subgroup prevalence in that population. The more refined the subgroup, the smaller *n*_*s*_ will tend to be and hence the resulting test will have lower power to detect a true effect. That is, rarer subgroups are harder to detect. Intuitively this highlights how the original trial design has reduced power to support more intricate subgroup discovery.

**Figure 1:**
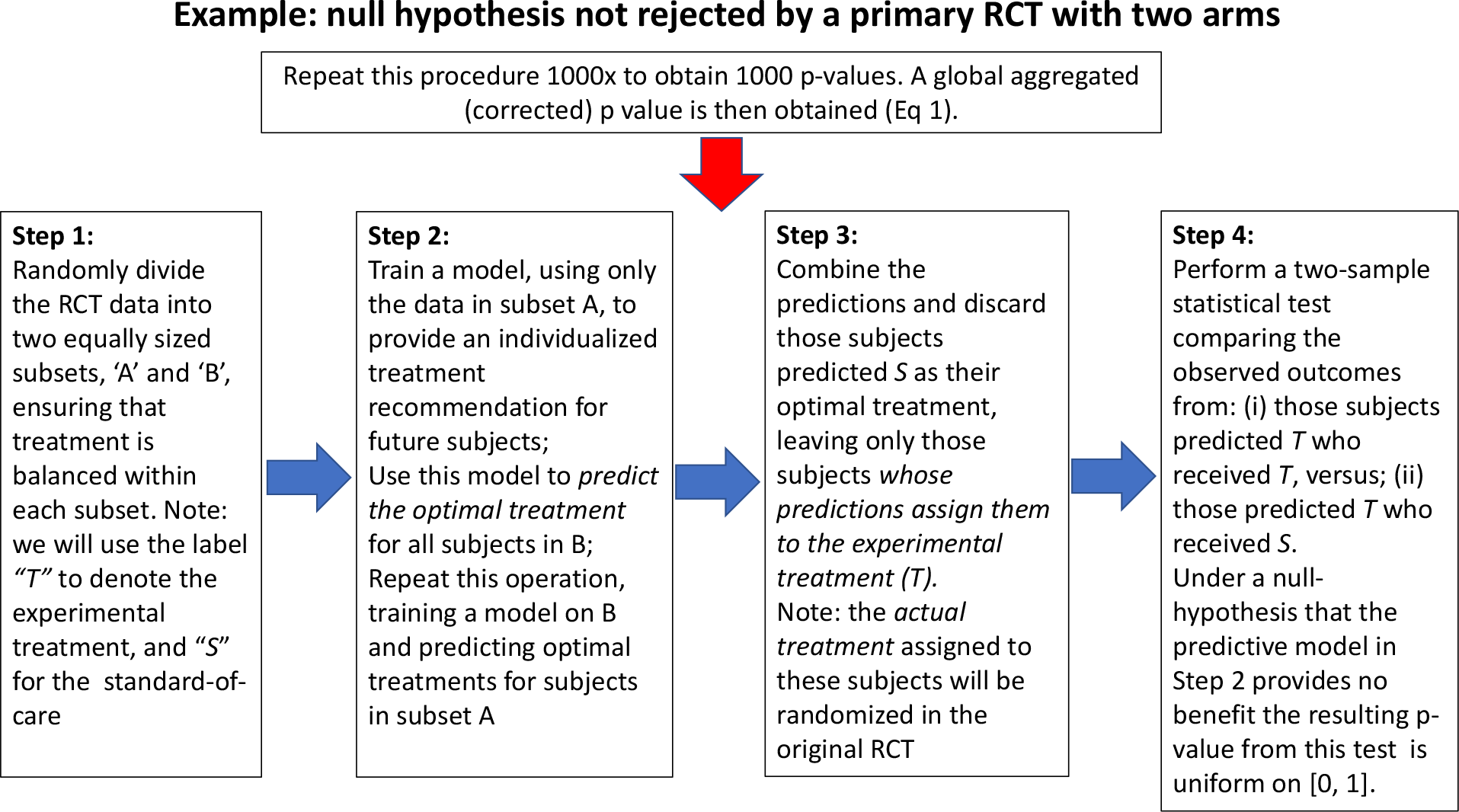
Illustrative example of hypothesis testing in exploratory subgroup discovery using 1000 iterations of two-fold cross-prediction. The example considers a primary RCT with two arms where a null hypothesis of “no improvement from the experimental treatment” is not rejected, i.e. no significant evidence of the experimental treatment providing improvement over the standard-of-care. Each random division results in a corresponding pvalue against the null hypothesis of no benefiting subgroup. The p-values are then aggregated for the overall test as given by Eq 1.

### False-positive control of non-crossovers: added predictive benefit of the ML analysis

The primary outcome in a standard RCT will often be strongly associated with particular baseline covariates and prognostic factors, e.g. severity of disease or co-morbidities. Generalised linear models (GLMs) provide one of most interpretable statistical models for relating clinical outcome to a combination of prognostic factors and the randomised treatment. Using more complex and therefore less interpretable ML methods needs to be justified with respect to the added benefit over this baseline model. In this context, the utility of ML methods is in their ability to detect non-linear interactions between prognostic factors and the randomised intervention. Using exactly the same data-splitting approach as for the discovery of statistically significant crossover subgroups, we can objectively evaluated the *added benefit* of the ML method. We illustrate the approach using a binary clinical outcome, *y*_*i*_ ∈ {0, 1} for the *i*’th subject, and a logistic regression GLM where

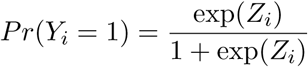

with linear predictor *Z*_*i*_ = *X*_*i*_*β* + *T*_*i*_*α*, for prognostic variables, *X*, and randomised treatment indicator *T*. The procedure is summarised below,

- For *K* iterations (e.g. *K* = 1000):

1. Split the data into two equally sized subsets with balanced number of treated and untreated cases in each half.
2. Fit a GLM to each subset separately and record for each individual their out-of-sample linear predictor 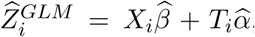 where 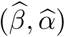 are obtained from the in-sample data fit.
3. Fit the ML method to each subset separately and predict the out-of-sample outcome probabilities, 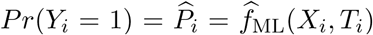, to obtain the corresponding log-odds out-of-sample prediction 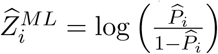 for each individual *i.*
4. Fit a ‘stacked’ GLM model to the full dataset using the *n ×* 2 matrix of prediction values 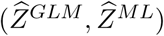 as two independent covariates variables,

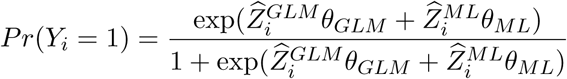

to obtain 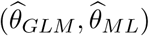. Record the p-value, *p*_*k*_, assigned to an ANOVA test for the model with *θ*_*ML*_ *≠* 0 versus a model with *θ*_*ML*_ = 0.
5. Construct the aggregate p-value from the set *p*_1_,…, *p*_*K*_ using adjustment method from Eq 1.

This method is analogous to ‘stacking’, a popular ML technique whereby multiple competing models are aggregated to form a more powerful ensemble model ^26^. We propose ‘stacking’ the standard accepted ‘vanilla’ statistical model (a GLM) alongside the predictions from a ML model. The added benefit of the ML based predictions can then be formally tested using Eq 1.

### Exploratory analysis

These ML driven procedures for testing both the presence of crossover and non-crossover TEH provide p-values uniformly distributed under the null. However, this approach is by definition non-constructive: the output does not contain an estimate of the discovered subgroup, nor an estimate of the TEH.

If the analysis leads to a rejection of the null (the aggregated p-value is below a pre-specified significance level), we then recommend fitting the same ML model to the full dataset and using this model to estimate the individual treatment effects. Further exploratory analysis can then be undertaken to assess the structure of the heterogeneity, such as: ‘which individuals are contained within the subgroup?’; ‘which covariates are predictive of TEH?’; ‘is the subgroup clinically relevant?’. This could be done via scatter plots of important covariates against the individual treatment effects. It is often possible to characterise a method detecting a true signal in the data by a few simple rules, for example using decision trees (e.g. Fig 2, panel D)^27^. By proceeding in this order, first evaluating the p-value for the null hypothesis, then undertaking the exploratory analysis using the full data, formal control of the type I error is still obtained.

**Figure 2:**
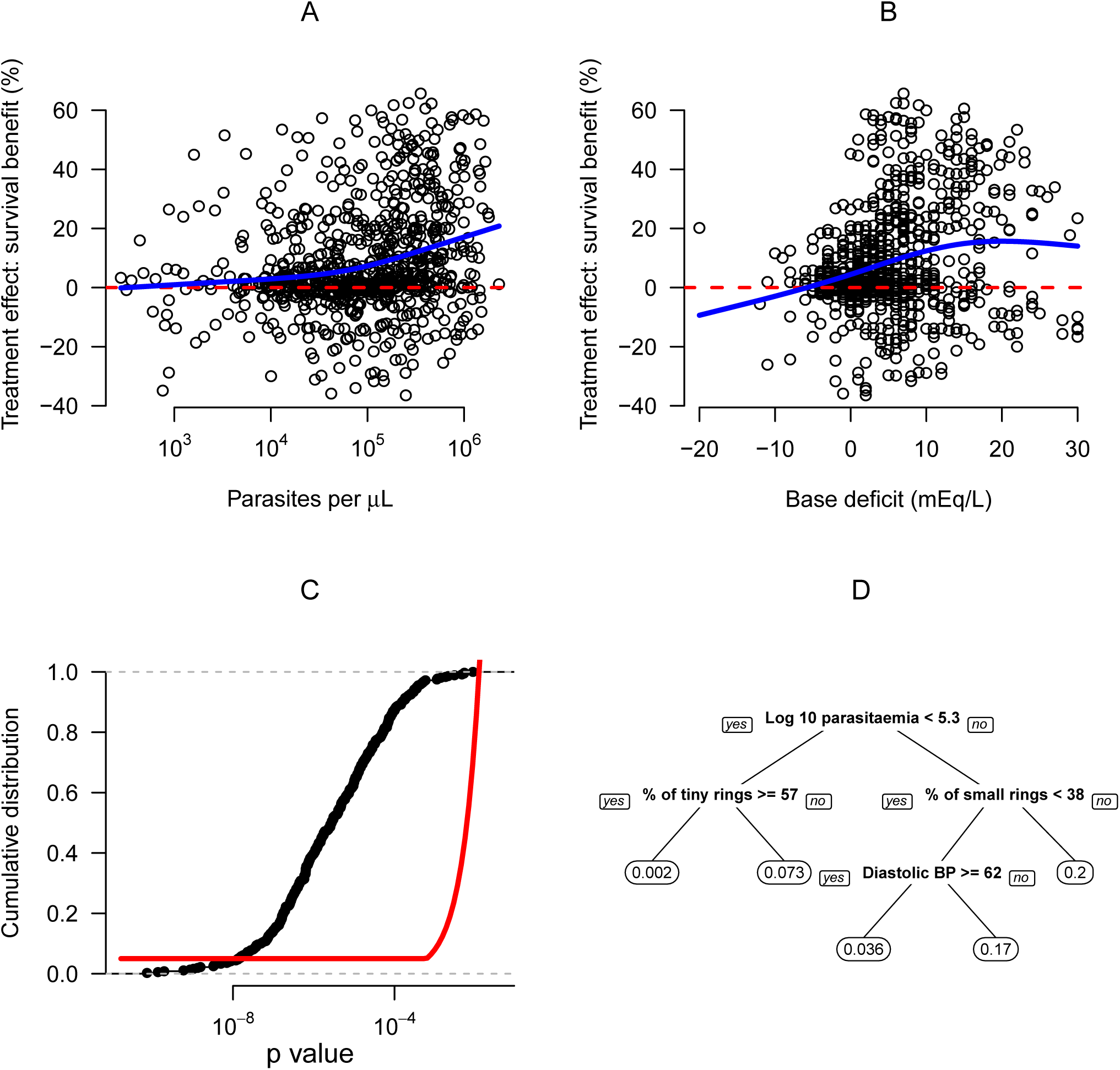
Graphical visualisation and validation of TEH defined by non-crossover interactions in the SEAQUAMAT trial. Panels A&B show the univariate relationships to the individual predicted treatment effect for total parasite biomass and base deficit, respectively. The thick blue lines show spline fits to the data. Panel C shows the cumulative distribution of the p-values for the added benefit of the ML model obtained by repeated data-splitting and stacking of the standard model alongside the ML model. Significance (at the 5% level) is obtained if the black line crosses above the red boundary (this is given by the function *Q*_*γ*_ in Eq 1). Panel D summarises the overall non-crossover interaction found by the RF model with a pruned regression tree model fitted to the individual treatment effects. The leaves of the tree in panel D show the mean treatment effect.

### Transparency and reproducibility

It is essential that all the findings and analysis paths taken are transparent and auditable to an external researcher. This can be achieved through the use of statistical notebooks, akin to the laboratory notebook in experimental science. Mainstream programming environments for data analysis (such as R and Python) provide open source notebooks such as R Markdown or Jupyter which seamlessly combine the analysis and the reporting. This allows all the exploratory analysis paths to be curated. Research recorded in a computational notebook is transparent, reproducible, and auditable. Auditability can be further improved without becoming burdensome through the use of version control repositories such as *github* (https://github.com) which record, timestamp, and preserve all versions and modifications of the analysis notebooks. Reproducibility can also be improved via the use of cloud computing services such as *Code Ocean*. In this way all of the steps, time lines, and historical evolution of the subgroup analysis are included and the work flow is open to external criticism and interrogation. Any published results can be audited back to the original RCT. Any p-values or statistical estimates that point toward subgroup effects that are reported subsequent to the heterogeneity tests need to be clearly labeled as such and treated with caution due to the potential for evidence inflation and post selective inference that arises from using the data twice. We prefer to label such measures that follow after data interrogation as qualitative, or q-values, as the formal statistical sampling uncertainty is often unknown ^28^.

## Results

### Antimalarial pharmacodynamics of artemisinin in severe malaria

Severe *Plasmodium falciparum* malaria is a medical emergency with case fatality rates ranging from 10 to 40% ^29^. A recent major advance in the treatment of severe malaria has been the introduction of parenteral artesunate. In Asia, this has been shown to reduce mortality by a third ^21^, and in Africa by a fifth ^30^. To illustrate the methodology advocated in this work, we use data from the definitive study of artesunate for severe malaria in Asia (SEAQUAMAT: South East Asian Quinine Artesunate Malaria Trial). This was a large multi-country randomised trial comparing intravenous quinine to intravenous artesunate^21^.

The superiority of parenteral artesunate for severe malaria is now well established ^31^. Thus in this retrospective analysis the artesunate arm is considered ‘standard-of-care’. The complete statistical analysis is published as an open source Code Ocean capsule and is entirely reproducible ^19^. This analysis provides an easily adjusted template for new exploratory subgroup analyses of different datasets.

We chose to use RFs to fit the data, one of the most popular and important ML methods in use today ^20^. This method deals well with multiple correlated covariates as is the case in these data. We first evaluate whether there is evidence for a crossover subgroup of patients who would benefit from quinine treatment as opposed to artesunate. The subgroup analysis does not reject the null hypothesis of “homogeneous optimal treatment allocation” (p = 1) showing that there is no evidence in the data of any subgroup benefitting from quinine.

This analysis was followed by examining the added benefit of the predictive RF model relating patient survival to the baseline measurements and treatment. An aggregation of the p-values obtained by repeated data-splitting and ‘stacking’ of the out-of-sample ML model predictions alongside the validated best linear predictor (the linear predictor on the logistic scale comprised of Glasgow coma scale, base deficit and treatment^32^) showed a strongly significant added benefit of the RF-ML model (p=10^*-*6^, Fig 2, panel C). Statistical significance of the repeated data-splitting and cross-prediction procedure can be assessed visually by comparing the cumulative distribution of the resulting p-values against the boundary curve as given by Eq 1.

Further exploratory analysis attempted to characterise possible interactions explaining this variation in predicted individual treatment effect. This analysis showed that significant TEH can be partially explained by the total non-sequestered parasite biomass (panel A) and the base deficit (panel B). This TEH can be summarised using a pruned CART model decision tree (panel D)^27^. This suggests that the greatest benefit of parenteral artesunate is seen in patients with large numbers of circulating young ring stage parasites (interaction between total parasitaemia and % of young rings). This is not highlighting a clinically relevant subgroup but helps elucidate the mechanism of action of artemisinin, a useful exercise in light of emerging drug resistance ^33^. Moreover, these results are concordant with the current proposed mechanism of action of the artemisinin derivatives and the importance of the artemisinin specific mode of action in the treatment of severe malaria. Artemisinins kill a broader range of parasite stages compared to quinine, notably the younger circulating ring forms, thereby reducing further sequestration and subsequent death in patients with a high parasite biomass ^34^.

## Discussion

We have shown how modern ML algorithms can be trained safely to determine the presence of TEH in a way that rigorously controls for type I error. The validity of our data splitting and out-of-sample prediction procedure holds irrespective of the method used, provided that samples are independently recruited from the study population. If this is not the case, for example if patients are recruited in pairs, or are related in some manner, then adjustments need to be made to ensure that the p-value reports the correct out-of-sample evidence. The choice of discovery algorithm should depend on the measurement variables collected (how many, and of which type) and the primary or secondary outcomes of the study for which TEH analysis is to be applied, e.g. survival time, binary outcome, continuous risk score. The specification of the stratifying measurements used by the method needs careful thought under a principle of ‘no free power’ in that feeding in irrelevant predictor variables will reduce the ability to detect true signals ^24^.

It is important that the analysis is transparent and the methods, data transformations, and analytic procedures are laid out and documented in an auditable plan, and that any code base used is properly documented and available for scrutiny. We recommend the use of open source repositories such as *github* or cloud computing services such as *Code Ocean* for fully reproducible data analyses. By following some simple guidelines we hope to improve upon the reliability and stability of TEH analysis reported in the literature.

Recent advances in statistical ML algorithms taken together with recent advances in measurement technologies have the potential to impact heavily and positively in the advancement of medical science. However alongside these advances great care must be taken to ensure that the integrity of the statistical analysis and the validity of the evidence base is upheld at all times.

### Panel 1: Overview of exploratory hypothesis generating ML-guided analysis.

- **TEH results in either *crossover* or *non-crossover* interactions** Crossover interactions imply that the optimal treatment allocation differs between patients (e.g. there is a subgroup of patients who’s optimal treatment is not the standard-of-care). Non-crossover interactions are important to understand intervention mechanisms and can be an important element in subsequent cost-benefit analyses.
- **Retrospective subgroup analysis** Before undertaking a retrospective hypothesis generating subgroup analysis on existing RCT data, it is necessary to write an ML subgroup statistical analysis plan (ML-SSAP) which should pre-specify the statistical or ML algorithm and the set of potential stratifying variables along with any potential explanatory, prognostic factors. This must define the ‘standard-of-care’ treatment (this could be different from when the trial was designed). More careful data preparation will increase the power to detect a true effect. The outcome variable should ideally match that used in the main trial.
- **Prospective subgroup analysis** We recommend including a ML-SSAP with the main trial protocol. In the same way as for a retrospective analysis, this must prespecify the variables included in the analysis and the algorithm used for the subgroup discovery. If the outcome variable is different from the main trial outcome, this should be explicit.
- **Cross-validation method for an unbiased assessment of subgroups** Aggregation of p-values from repeated two-fold data-splitting can provide an unbiased p-value relating to the null hypothesis of ‘no crossover TEH’. This p-value can be taken at face value and if below a pre-specified significance level, the proposed subgroups from a full data analysis (fitting the same model to the full dataset) can be used to inform further trials.
- **Further exploratory analyses** As data are accrued and analysed, further reactive analyses may be of interest. Such exploratory analysis is recommended but should be clearly distinguished from the main pre-specified subgroup analysis. The p-values generated from these analyses can be denoted ‘q-values’ (qualitative p-values).
- **Statistical notebooks** The entire subgroup discovery process should be undertaken using computational notebooks (e.g. RMarkdown, Jupyter). Combined with version control tools such as *github* and cloud computing such as *Code Ocean* this allows for a fully reproducible and transparent process.

## Acknowledgments

We thank the Mahidol Oxford Research Unit for providing us with the data from the SEAQUAMAT trial. We are grateful to Nicholas White and Stije Leopold for valuable input concerning the interpretation of the SEAQUAMAT analysis. We thank Aimee Taylor for help with the graphical representation of the method. We thank Dr Rajen Shah for pointing us to the work by Meinshausen *et al.* on p-value aggregation.

## Author Contributions

The authors contributed equally to all parts of this work.

Abbreviations

RCTs: randomised controlled trials
ML: machine learning
TEH: treatment effect heterogeneity
RF: random forests
ML-SSAP: machine learning based subgroup statistical analysis plan
GLM: generalised linear model

